# Mitochondrial calcium modulates odor-mediated behavioral plasticity in *C. elegans*

**DOI:** 10.1101/2023.08.29.555059

**Authors:** Hee Kyung Lee, Dong-Kyu Joo, Kyu-Sang Park, Kyoung-hye Yoon

**Author notes:** Department of Cell, Developmental and Integrative Biology, University of Alabama at Birmingham, Birmingham, AL 35233, USA. Co-corresponding authors (KSP) (KHY).

## Abstract

Despite growing understanding of the various roles mitochondria play in neurons, how they contribute to higher brain functions such as learning and memory remains underexplored. Here, using the nematode *Caenorhabditis elegans,* we found that the mitochondrial calcium uniporter (MCU) pore forming unit MCU-1 is required for aversive learning to specific odors sensed by a single sensory neuron, AWC^ON^. MCU-1 expression was required in the sensory neuron at the time of odor conditioning for proper behavioral response to 60 min of prolonged odor exposure. Through genetic and pharmacological manipulation, we show evidence that MCU is activated in response to prolonged odor conditioning, causing mtROS production, leading to NLP-1 secretion. Finally, we show that the timing of MCU activation and neuropeptide release correspond with the OFF-neuron properties of the AWC neuron, suggesting that mitochondrial calcium entry and neuropeptide secretion coincide with AWC activation upon odor removal. Overall, our results demonstrate that, by regulating mitochondrial calcium influx, mitochondria can modulate the synaptic response to incoming stimuli in the sensory neuron, resulting in learning and modified behavior.

## Introduction

Learning and memory allow past experience of an organism to adjust and shape their behavior, thereby increasing its fitness and chance of survival. This behavioral plasticity is modulated by multiple factors such as the length and frequency of the experience, or concurrent external and internal sensory inputs. Such set of factors ensures that the learning and strength of the memory is commensurate to the type of experience and its context.

The cellular and molecular substrate for learning and memory is the change in synaptic activity strength. Among the numerous factors involved in modulating synaptic activity, mitochondria are reported to play a role by supplying ATP through energy metabolism(Ashrafi *et al*, 2020; Hebert-Chatelain *et al*, 2016; Rangaraju *et al*, 2014; Verstreken *et al*, 2005), buffering calcium(Alnaes & Rahamimoff, 1975; David *et al*, 1998; Lee *et al*, 2007; Tang & Zucker, 1997), and producing localized ROS(Jia & Sieburth, 2021; Zhao *et al*, 2018). Consistent with this, a few studies have shown instances where mitochondria are involved in learning and memory(Duan *et al*, 2021; Hebert-Chatelain *et al*., 2016; Oettinghaus *et al*, 2016), and mitochondrial dysfunction in the brain has been linked to synaptic dysfunction and decline in cognitive abilities(Todorova & Blokland, 2017). Further studies of mitochondria’s role in neuronal function are needed to understand the various ways in which metabolic activities and their dysfunction are interconnected with higher brain functions such as learning and memory.

The mitochondrial calcium uniporter (MCU) complex is the main route of calcium entry into mitochondria. The complex consists of MCU channel protein and several regulatory subunit proteins that play modulatory roles. In *C. elegans*, the complex consists of the channel protein MCU-1, along with two regulatory subunit proteins MICU-1 and EMRE-1. With the molecular identity revealed a little more than a decade ago(Baughman *et al*, 2011; De Stefani *et al*, 2011), studies are beginning to uncover its various roles in different tissues during normal and pathological conditions (Burkewitz *et al*, 2020; Higashitani *et al*, 2023). Even so, the mechanisms by which the channel is modulated in various tissues under physiological conditions remain poorly understood, and its functions and regulation in neurons are still emerging areas of study.

The nematode *C. elegans* displays complex behavior that can be modified through past experience. Its simple nervous system consisting of 302 neurons, its genetic tractability, and the array of tools available, offer the chance to investigate the detailed molecular underpinnings of learning and memory. *C. elegans* can learn to increase or decrease its attraction to odors based on past experience, and can control the degree of these modifications depending on the context and length of the experience(Colbert & Bargmann, 1995; Kauffman *et al*, 2010; Lee *et al*, 2010). For attractive odors such as butanone and benzaldehyde that are mainly detected by the chemosensory neuron AWC, prior experience to the odor in the absence of food makes them less attractive to the worms. How much less, is dependent on the length of the experience, and a gradual decline in attraction can be seen from 0 to 2 hours of prior exposure (Lee *et al*., 2010). Interestingly, this progressive decline is mediated by distinct molecular steps and can be genetically separated. For example, *cng-3* in AWC is required for the mild decrease in attraction after 30 min of odor conditioning (O’Halloran *et al*, 2017). However, aversive learning to 60 min conditioning is unaffected in *cng-3* mutants, demonstrating the distinct molecular steps involved to produce the seemingly smooth continual decline in attraction. Similarly, nuclear translocation of the cGMP-dependent protein kinase EGL-4 in AWC is required for aversive learning to 60 min or longer conditioning (Lee *et al*., 2010). Thus, the aversive odor learning paradigm in *C. elegans* offers a compelling system to study the molecular mechanisms of behavioral plasticity.

In this study, we use *C. elegans* to show that MCU facilitates odor learning, and thus change in odor behavior, by controlling neuropeptide release from the primary sensory neuron. We found that *mcu-1* mutants are defective for aversive learning to AWC^ON^-sensed odors. MCU-1 was required in the AWC neuron, at the time of learning. The learning defect was specific to learning after 60-min conditioning, which requires the NLP-1 neuropeptide. By genetic and pharmacological manipulation, we show that MCU in the AWC responds to 60 min odor exposure to produce mitochondrial ROS(mtROS), which triggers NLP-1 secretion from the same neuron. Moreover, ectopic activation of MCU or mtROS production during odor exposure was sufficient to accelerate learning. Lastly, blocking MCU-1 at different time points during the 60 min conditioning revealed that MCU-1 was needed towards the end of the prolonged conditioning, which suggests mitochondrial calcium influx, and thus neuropeptide release, is coupled to the AWC calcium transient that occurs upon odor removal. Taken together, our study shows that mitochondria calcium in neurons contribute to learning by modulating the synaptic response to sensory stimuli.

## Results

### *mcu-1* mutants display AWC^ON^-specific aversive learning defect

To study the role of MCU in neurons, we tested a null mutant strain of *mcu-1* for behavioral defects, including odor behaviors. *C. elegans* detects attractive volatile odors mainly through two pairs of sensory neurons, AWC and AWA. The AWC neuron pair is asymmetric, and expresses different sets of receptors(Wes & Bargmann, 2001). With this in mind, we selected odorants that represented the function of each sensory neurons: diacetyl, 2-butanone, 2,3-pentanedione, and benzaldehyde, which are mainly sensed by AWA, AWC^ON^, AWC^OFF^, both AWCs, respectively (Fig 1A). Mutant worms showed normal chemotaxis for all odorants tested, indicating that worms can sense and move toward attractive odors in the absence of MCU-1 (Fig 1B-C).

**Fig 1.**
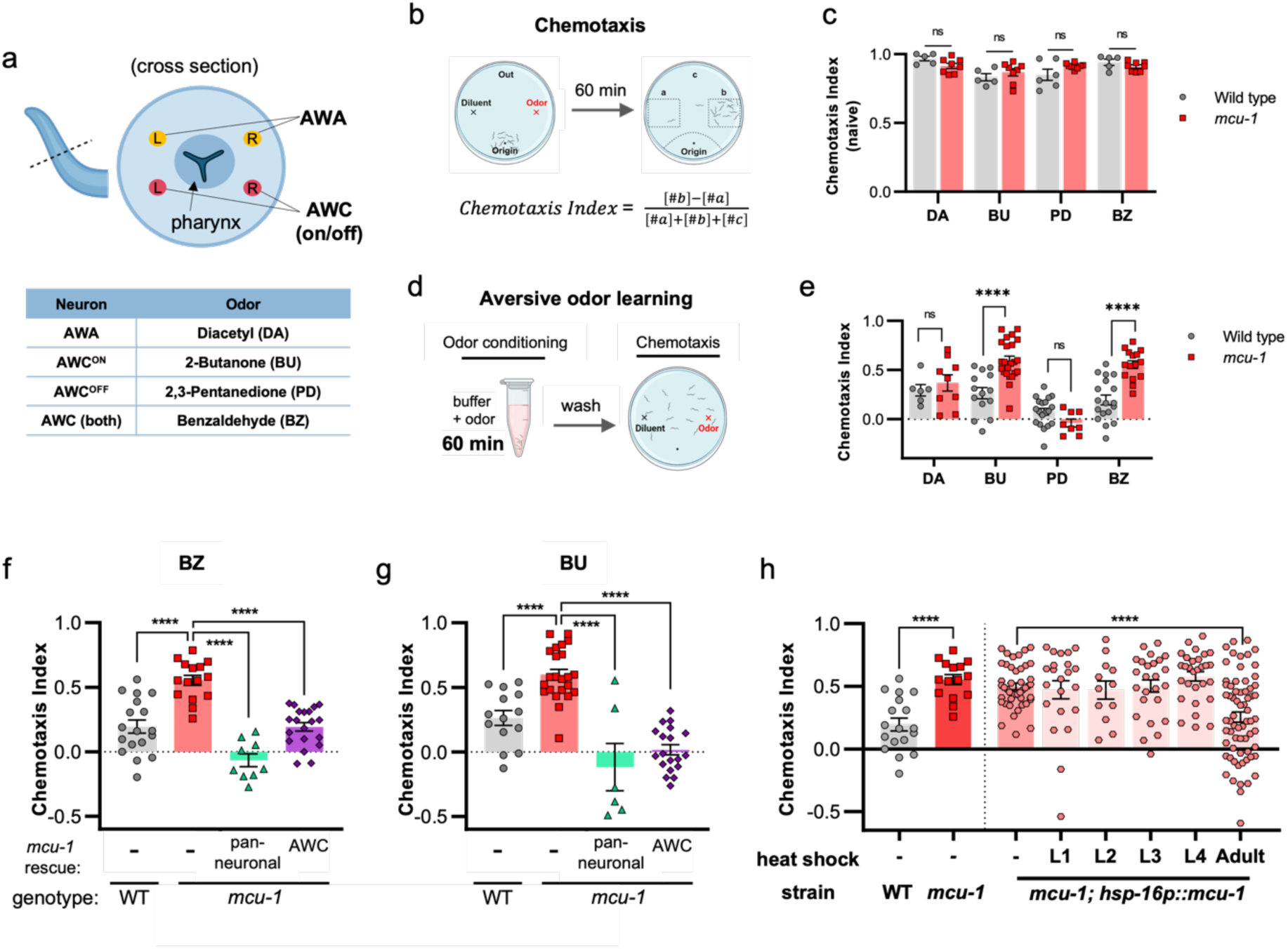
*mcu-1* is required in the AWC^ON^ sensory neuron at the adult stage for aversive odor learning. (A) Each *C. elegans* chemosensory neuron pairs detect distinct attractive odorants. (B) Schematic for chemotaxis assay and chemotaxis index (CI) calculation. (C) Chemotaxis index for naïve N2 and *mcu-1(tm6026)* worms. (D) Schematic for aversive odor learning. (E) Chemotaxis index in worms after 60 min odor conditioning. (F,G) Aversive odor learning in neuron(*rab-3p*)- and AWC(*ceh36p*)-specific rescue strains for benzaldehyde(BZ) and butanone(BU). (H) Chemotaxis index of transgenic strains expressing MCU-1 under the heat-shock-inducible promoter (*hsp-16.2p*). Worms were given heat shock at different larval stages and tested as day 1 adults. P values were determined using Student’s t-test (C, E, comparison between WT and *mcu-*1 in H) or one-way ANOVA with Dunnett’s multiple comparisons test (F, G, comparison between larval-stage-heatshock in H). Asterisks indicate p value (ns, not significant; ****p<0.0001).

We next tested *mcu-1* mutants for odor learning behaviors. In the aversive odor learning paradigm, previous exposure to odors without food causes worms to ignore or avoid what used to be an attractive odor (Figure 1D). After 60 min of pre-exposure, *mcu-1* mutants failed to display aversive learning to benzaldehyde and butanone, whereas learning for other odors remained normal (Fig 1E). Since butanone is sensed by AWC^ON^, and benzaldehyde is sensed by both AWC neurons, this suggested that learning was specifically defective for AWC^ON^. Consistent with this, pentanedione, which is sensed by the AWC_OFF_ neuron, showed normal aversive learning. Thus, *mcu-1* mutants fail to learn odors sensed by the AWC^ON^ neuron.

### MCU-1 is required in the primary sensory neuron for aversive odor learning

We next sought to find where MCU-1 was needed for odor learning. We hypothesized that MCU-1 is likely needed in the neurons. To test this, we generated a transgenic strain that expressed MCU-1 under a neuron-specific promoter (*rab-3p*). As expected, neuronal *mcu-1* rescue strain showed normal aversive learning to both butanone and benzaldehyde (Fig 1F,G). To further narrow down the neuron responsible, we next made a cell-specific *mcu-1* transgenic rescue strain. We hypothesized that MCU-1 is likely needed in the neuronal circuit that includes the primary sensory neuron AWC^ON^ and a few immediately downstream interneurons (Chen *et al*, 2006; White *et al*, 1986). When we expressed MCU-1 under the AWC-specific promoter (*ceh-36p*), we found that it was also sufficient to fully restore aversive learning (Fig 1F,G). Thus, MCU-1 is needed in the sensory neuron AWC to mediate odor learning.

### MCU-1 is required post-developmentally

We next wondered when MCU-1 needed for odor learning. *C. elegans* goes through 4 larval stages until they reach adulthood. For most neurons, cell fates are determined during the embryonic stage and neurons are fully functional by the time of hatching . However, strength of neuronal response may change as the worm grows and matures (Fujiwara *et al*, 2016). To determine at which stsage MCU-1 is needed for odor learning, we generated a transgenic strain that expressed MCU-1 under a heat-inducible promoter. Inducing MCU-1 expression at different larval stages revealed that aversive learning was restored only when MCU-1 was induced at the adult stage, which was only a few hours before they were used for the aversive learning assay (Fig 1H). This showed that MCU-1 is required post-developmentally, and close to the time of learning. We also noted that the ineffectiveness of earlier MCU-1 induction may speak to the rate of turnover of MCU-1, and the need for continuous supply of newly synthesized proteins (Fig 1H). In addition, imaging fluorescently labeled AWC neurons revealed no change in morphology from wild type to mutant (Fig S1), adding to the evidence that the role of MCU-1 in odor learning is post-developmental. Thus, MCU-1 is required at the adult stage for odor learning.

### Aversive learning defect in *mcu-1* is specific to 60 min conditioning

In aversive odor learning, attractiveness to an odor progressively declines with longer pre-exposure, or conditioning, time (Lee *et al*., 2010). For AWC-mediated odor learning, some of these steps are molecularly distinct: learning to 30 min conditioning requires *cng-3*(O’Halloran *et al*., 2017), and further decrease in CI after 60 min or more requires translocation of the cGMP-dependent protein kinase EGL-4 into the nucleus(Lee *et al*., 2010).

To further characterize the learning defect of *mcu-1* mutants, we tested different conditioning times ranging from 30 to 120 min (Figure 2A). At 30 min conditioning, both wild type and *mcu-1* mutant strain showed a modest decrease in attraction (Figure 2B,C). At 60 min pre-exposure, wild type showed further decrease in CI, whereas *mcu-1* mutants remained at the 30 min level. However, after 90 min and beyond, both wild type and *mcu-1* worms similarly displayed a much lower CI close to 0. The pattern was observed for both butanone and benzaldehyde (Figure 2B,C). Although 90 min exposure to benzaldehyde resulted in slightly lower CI in *mcu-1* mutants than in wild type, in other trials this difference was not as pronounced (Figure 2D). Thus, to our surprise, the learning defect was specific to 60 min conditioning, but dispensable for shorter and longer conditioning times.

**Fig 2.**
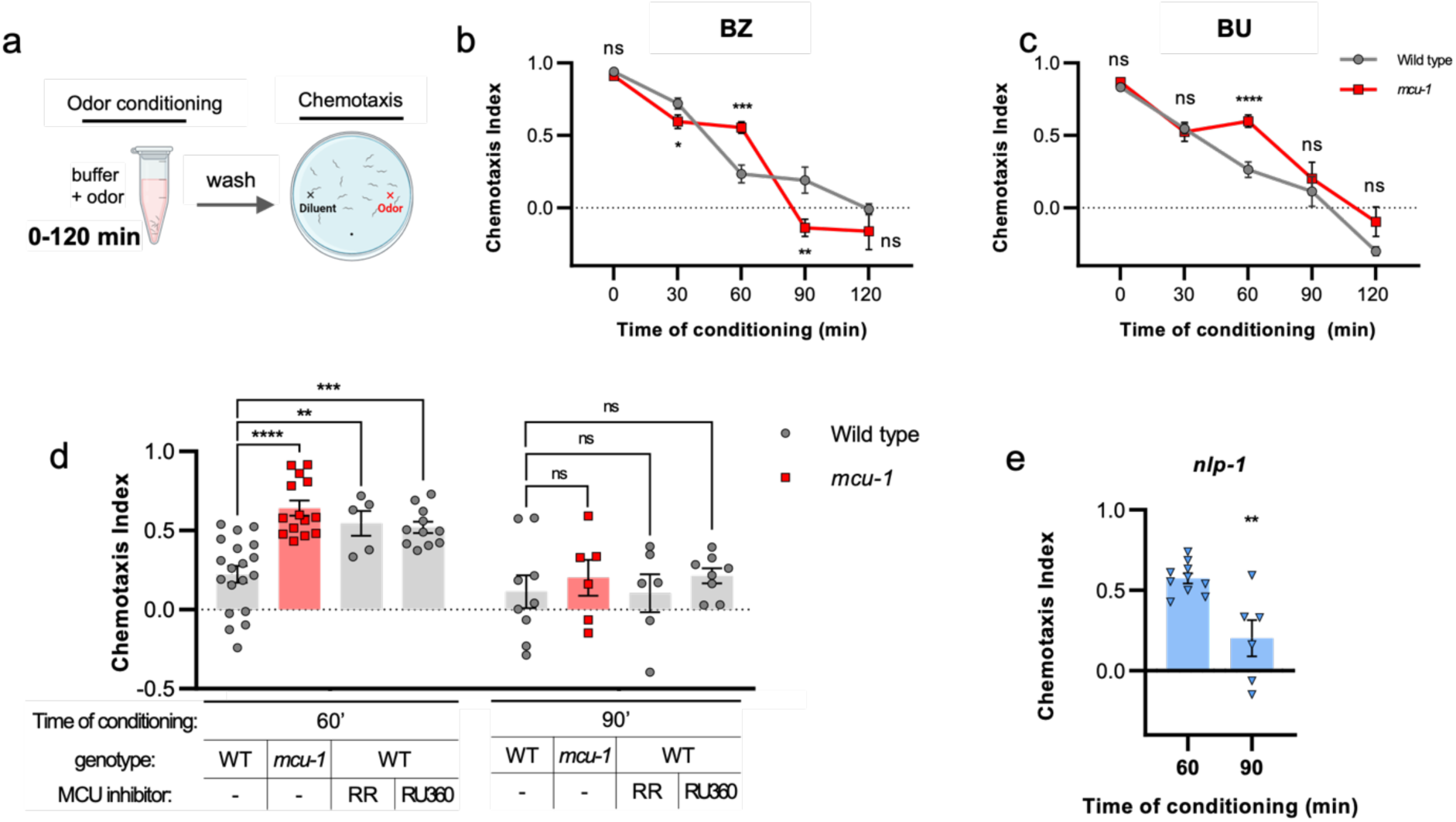
Mutants of *mcu-1* are specifically defective for 60 min odor learning. (A) Aversive odor learning was conducted with varied conditioning times. (B,C) Chemotaxis index after different conditioning times (ný6 trials per group). Statistics were (D) Wild type worms treated with pharmacological blockers of MCU, ruthenium red(RR) and RU360, display the same learning defect to butanone as *mcu-1* mutants. (E) *nlp-1* mutant is also defective for aversive learning after 60 min conditioning, but not after 90 min. P values were determined using Student’s t-test (B,C,E) or one-way ANOVA with Dunnett’s multiple comparisons test (D). Asterisks indicate p value (ns, not significant; *p<0.05; **p<0.01; ***p<0.001; ****p<0.0001).

### MCU function is required at the time of conditioning

Earlier, we found that MCU-1 expression is required post-developmentally for odor learning. To further narrow down when MCU-1 function is needed, we used ruthenium red and RU360, pharmacological blockers of MCU(Ying *et al*, 1991). When added to the odor solution during conditioning, we found that both ruthenium red and RU360 mimicked the *mcu-1* mutant phenotype: treatment during 60 min pre-exposure inhibited odor learning, but treatment during 90 min pre-exposure yielded normal odor learning indistinguishable from wild type (Fig 2D). Thus, pharmacological blockers of MCU can be used to reproduce *mcu-1* mutant phenotype, and MCU-1 function is needed at the time of conditioning for successful odor learning.

### MCU mediates NLP-1 neuropeptide secretion from AWC

While it was surprising that MCU-1 was required for learning to a specific length of conditioning time, a similar learning defect had been described previously for another mutant. Chalasani *et al*. had reported that mutant of a neuropeptide gene *nlp-1* showed defective aversive learning for isoamylalcohol, another AWC-sensed odor(Chalasani *et al*, 2010). Interestingly, the learning defect was also specific for 60 min conditioning, as mutants showed normal learning for 90 min conditioning(Chalasani *et al*., 2010). NLP-1 is expressed in many neurons, including AWC(Nathoo *et al*, 2001; Taylor *et al*, 2021). Conducting the same aversive learning experiment on *nlp-1* mutants, but with butanone, confirmed their results (Fig 2D). This suggested that *mcu-1* and *nlp-1* may be acting through the same pathway.

One likely possibility was that MCU-1 in the AWC is involved in NLP-1 secretion from the same neuron. To test this, we generated a transgenic strain that expressed Venus-tagged NLP-1 in the AWC neurons. The fluorescently tagged NLP-1 is not functional, but it is secreted into the body cavity, accumulating in the scavenging coelomocytes, whose fluorescence can be used as a proxy for neuropeptide secretion (Figure 3A)(Sieburth *et al*, 2007).

**Fig 3.**
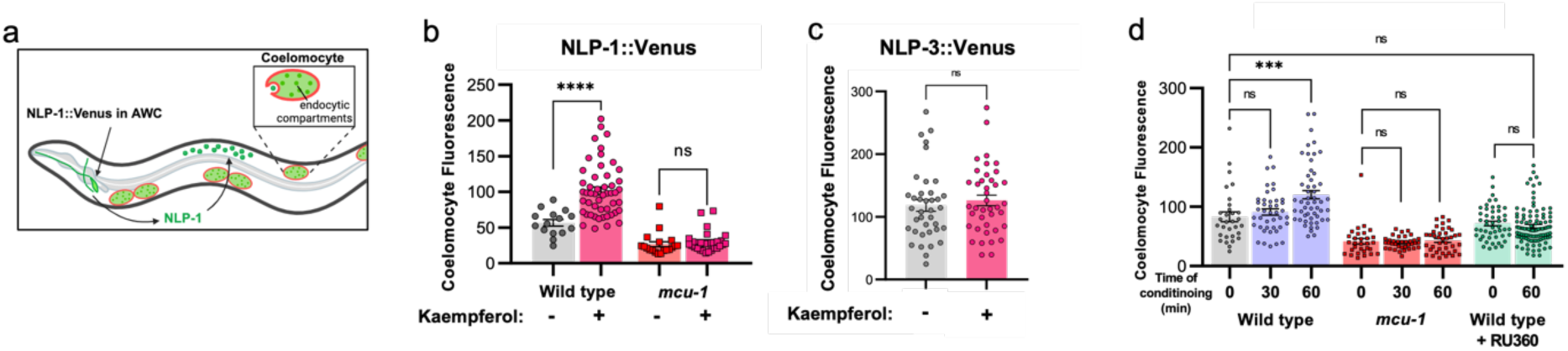
MCU activation or butanone exposure causes NLP-1 secretion from AWC neurons. (A) Diagram of NLP-1 secretion from AWC and uptake by coelomocytes. (B) NLP-1::Venus fluorescence measured in coelomocytes with and without 10 min treatment with the MCU activator kaempferol. (C) NLP-3::Venus fluorescence in coelomocytes with and without 10 min kaempferol treatment. (D) NLP-1::Venus fluorescence in coelomocytes after butanone conditioning. P values were determined using Student’s t-test (B,C) or one-way ANOVA with Dunnett’s multiple comparisons test (D). Asterisks indicate p value (ns, not significant; ***p<0.001; ****p<0.0001).

We first tested whether activating MCU causes NLP-1 secretion. For this, we treated wild-type worms with a MCU activator, kaempferol (Montero *et al*, 2004). After 10 min treatment with kaempferol, we found a significant increase in the fluorescent signal in the coelomocytes (Fig 3B). Kaempferol treatment in *mcu-1* mutants, as expected, did not result in fluorescence increase, showing that activation of MCU, and thus calcium influx into the mitochondria, is required for NLP-1 release (Fig 3B). Interestingly, MCU activation does not result in increased secretion of all neuropeptides from AWC, as no change in fluorescence was observed when another highly expressed neuropeptide, NLP-3 secretion, was monitored (Fig 3C).

### Prolonged odor exposure leads to NLP-1 secretion from AWC

If ectopic activation of MCU results in NLP-1 secretion, can we observe the same in response to a natural stimulus, such as 60 min odor conditioning? To find out, we exposed the NLP-1::Venus transgenic strain to butanone for either 30 or 60 min. We found that wild-type worms exposed to butanone for 60 min showed increased fluorescence in the coelomocyte, whereas no increase was observed after 30 min exposure (Fig 3D). This indicates that NLP-1 is secreted only after prolonged odor exposure, and 30 min is insufficient. When we tested *mcu-1* mutant worms, they did not show any increase in fluorescence in response to butanone regardless of exposure time (Fig 3D). These results were all consistent with the learning defect of *nlp-1* and *mcu-1* mutants, which are defective only for aversive learning to 60 min conditioning. Does NLP-1 secretion in response to prolonged odor conditioning require MCU function? To find out, we treated NLP-1::Venus worms with the MCU blocker RU360 during 60 min odor conditioning. In the presence of the MCU blocker, there was no increase in fluorescence in the coelomocytes (Figure 3D). Taken together, our results show that odor exposure of 60 min, but not 30 min, causes secretion of NLP-1 from AWC, in an MCU-dependent manner.

### mtROS is required for NLP-1 secretion and odor learning

How does MCU-1 mediate NLP-1 secretion? Calcium influx into the mitochondria stimulates ATP production as well as ROS production, both of which can support synaptic signaling(Jia & Sieburth, 2021; Rangaraju *et al*., 2014). Since a previous *C. elegans* study had shown that *mcu-1* was required for mtROS-mediated neuropeptide secretion from a pair of interneurons, we wondered whether NLP-1 secretion from the AWC sensory neuron was occurring through a similar mechanism(Jia & Sieburth, 2021).

To directly verify whether 60 min butanone exposure results in ROS production at the mitochondria, we generated a transgenic strain that expressed the mitochondrially targeted fluorescent ROS sensor mito-roGFP in the AWC neurons. We exposed the transgenic strain with butanone for 60 min and assessed the redox ratio of roGFP. We also exposed the strain to H_2_O_2_ for 10 min as positive control. We then assessed whether increased oxidation could be seen after each treatment. We specifically looked at mitochondria in the axons, since axonal mitochondria is known to be responsible for synaptic signaling and neuropeptide secretion (Jia & Sieburth, 2021). In both conditions, mitochondria in the AWC axons indeed showed increased oxidation (Figure 4A). Thus, prolonged odor stimulus causes ROS generation in the mitochondria of the sensory neuron.

**Fig 4.**
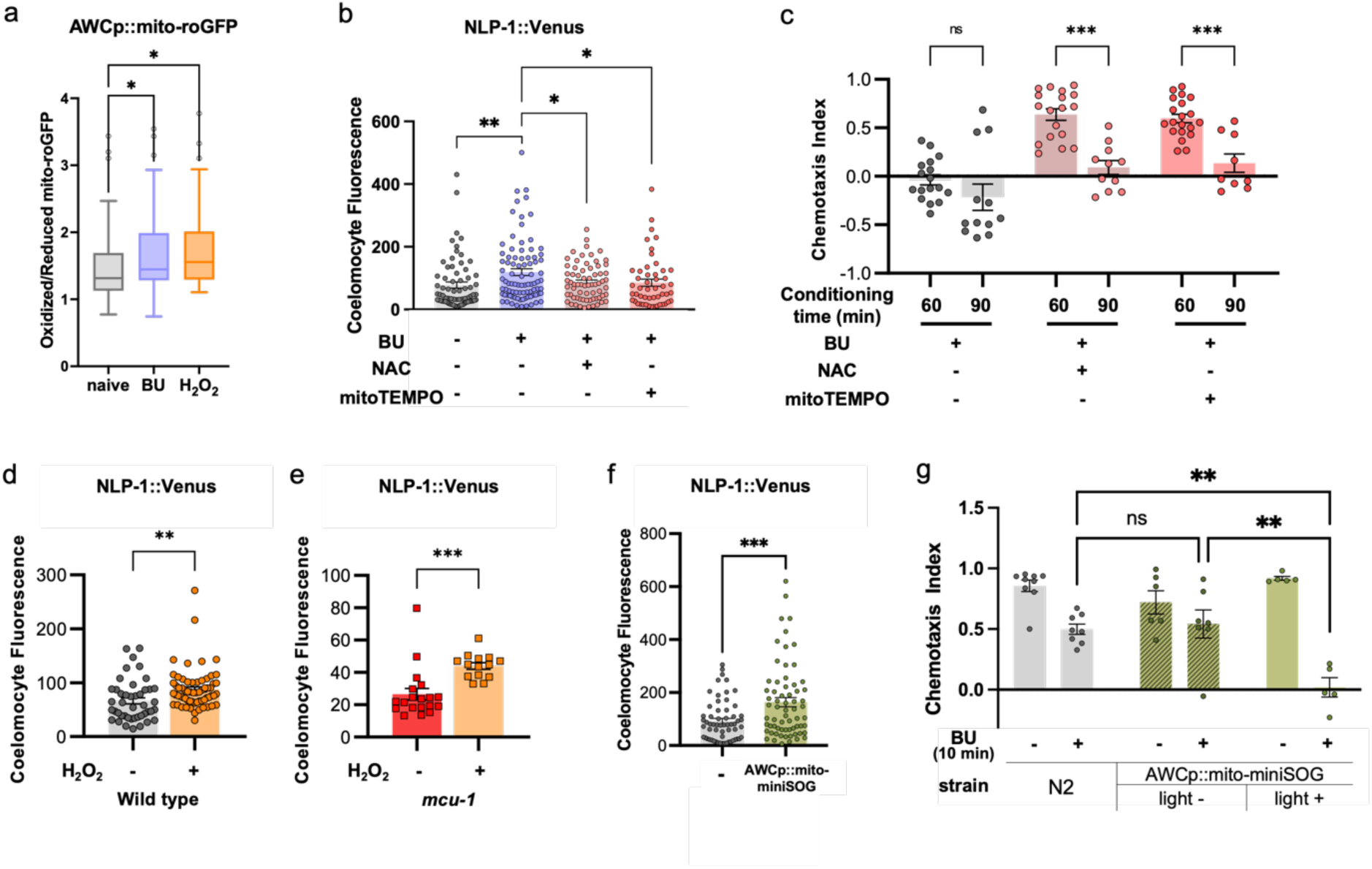
Odor conditioning causes mtROS production in AWC. (A) roGFP localized to AWC mitochondria was used to determine the redox ratio after exposure to 10 min H_2_O_2_ or 60 min butanone. (B) NLP-1 secretion is prevented by treatment with the antioxidants NAC and mitoTEMPO during odor conditioning. (C) Antioxidant treatment inhibits 60 min aversive odor learning, but not 90 min learning. (D,E) 10 min treatment with H_2_O_2_ is sufficient for NLP-1 secretion in both wild type and *mcu-1* mutant. (F) Expression of mitochondrially targeted miniSOG in the AWC neuron is sufficient for NLP-1 release under ambient light. (G) Change in chemotaxis index after 10 min odor conditioning. Activation of miniSOG by light causes fast odor learning. P values were determined using Student’s t-test (C,D,E,F) or one-way ANOVA with Dunnett’s multiple comparisons test (A,B,G). Asterisks indicate p value (ns, not significant; *p<0.05; **p<0.01; ***p<0.001).

Next, we reasoned that if mtROS is required for NLP-1 secretion, then treating with antioxidants that neutralize ROS should prevent its secretion. We therefore treated worms with N-acyl cysteine (NAC), a general antioxidant, or mitoTEMPO, which specifically scavenges mitochondrial ROS, during odor conditioning. Consistent with our hypothesis, worms treated with either NAC or mitoTEMPO failed to show NLP-1::Venus accumulation in the coelomocytes (Figure 4B). Consistently, NAC and mitoTEMPO were also effective in preventing odor learning at 60 min, whereas it had no effect on 90 min learning, exactly matching the *mcu-1* mutant phenotype (Figure 4C). Thus, mtROS is required for NLP-1 secretion, as well as 60-min odor learning.

Superoxide anions are the most abundant ROS produced by the mitochondria and is quickly converted into H_2_O_2_ by superoxide dismutase (SOD)(Zorov *et al*, 2014). H_2_O_2_ can alter activities of target enzymes by modifying the thiols of cysteine residues(Veal *et al*, 2007). Therefore, we next asked whether adding H_2_O_2_ or inducing cell-specific mtROS would cause NLP-1 secretion without the prolonged odor exposure. To do this, NLP-1::Venus worms were treated with 5 mM H_2_O_2_ for 10 min and coelomocyte fluorescence was measured. We found that 10 min H_2_O_2_ treatment was sufficient to induce NLP-1 secretion (Figure 4D). Moreover, H_2_O_2_ treatment of the *mcu-1* mutant was also sufficient to induce NLP-1 secretion (Figure 4E). This contrasts with the earlier kaempferol treatment, which was not effective in *mcu-1* mutants (Figure 3B). This shows that H_2_O_2_ production occurs downstream of MCU activation and calcium influx into the mitochondria.

To achieve precise mtROS generation only in the AWC neuron, we expressed the light-activated singlet oxygen generator, miniSOG, in the AWC mitochondria. When illuminated by blue light, ROS produced by miniSOG causes cell death in the cells that express it, making it a useful tool for cell ablation. However, shorter illumination can be used for ROS generation without causing cell ablation(Jia & Sieburth, 2021). When the transgenic strain was crossed with the nlp-1::Venus strain, we found that the double transgenic strain showed markedly higher Venus fluorescence in the coelomocytes (Figure 4F). Thus, mtROS in AWC can induce NLP-1 secretion from the same neuron.

### Coincidental activation of MCU and odor exposure accelerates odor learning

Our data so far suggest that prolonged exposure to AWC^ON^-specific odors cause MCU activation, causing calcium influx and mtROS production, which is required for 60-min odor learning. Based on this, we hypothesized that if we ectopically induce mtROS, odor learning can happen with only a brief exposure to odor. Normally, 10 min of butanone conditioning results in only a slight decrease in CI (Fig 4G). However, we found that the same treatment in worms expressing mito-miniSOG in AWC neurons produced a sharp decline in CI that is close to zero. This was dependent on light-activated mtROS generation by miniSOG, as the same procedure conducted without light illumination showed CI similar to that of wild type (Fig 4G). Also, mito-miniSOG worms conditioned in buffer without odor showed normal chemotaxis, demonstrating that odor must be presented simultaneously with mtROS for learning to occur.

Similarly, simultaneous activation of MCU with odor exposure also accelerated odor learning. Treating worms with kaempferol alone for 10 min resulted in high CI identical to naïve worms, showing that MCU activation and NLP-1 secretion alone is not sufficient to produce decreased attraction to odors. However, when kaempferol was added together with butanone for 10 min, worms exhibited a far lower CI that is typical of 60-min conditioning (Fig 5A). Taken together, when odor stimulus is paired with MCU activation, which produces mtROS, which in turn causes NLP-1 secretion, aversive odor learning is accelerated.

**Fig 5.**
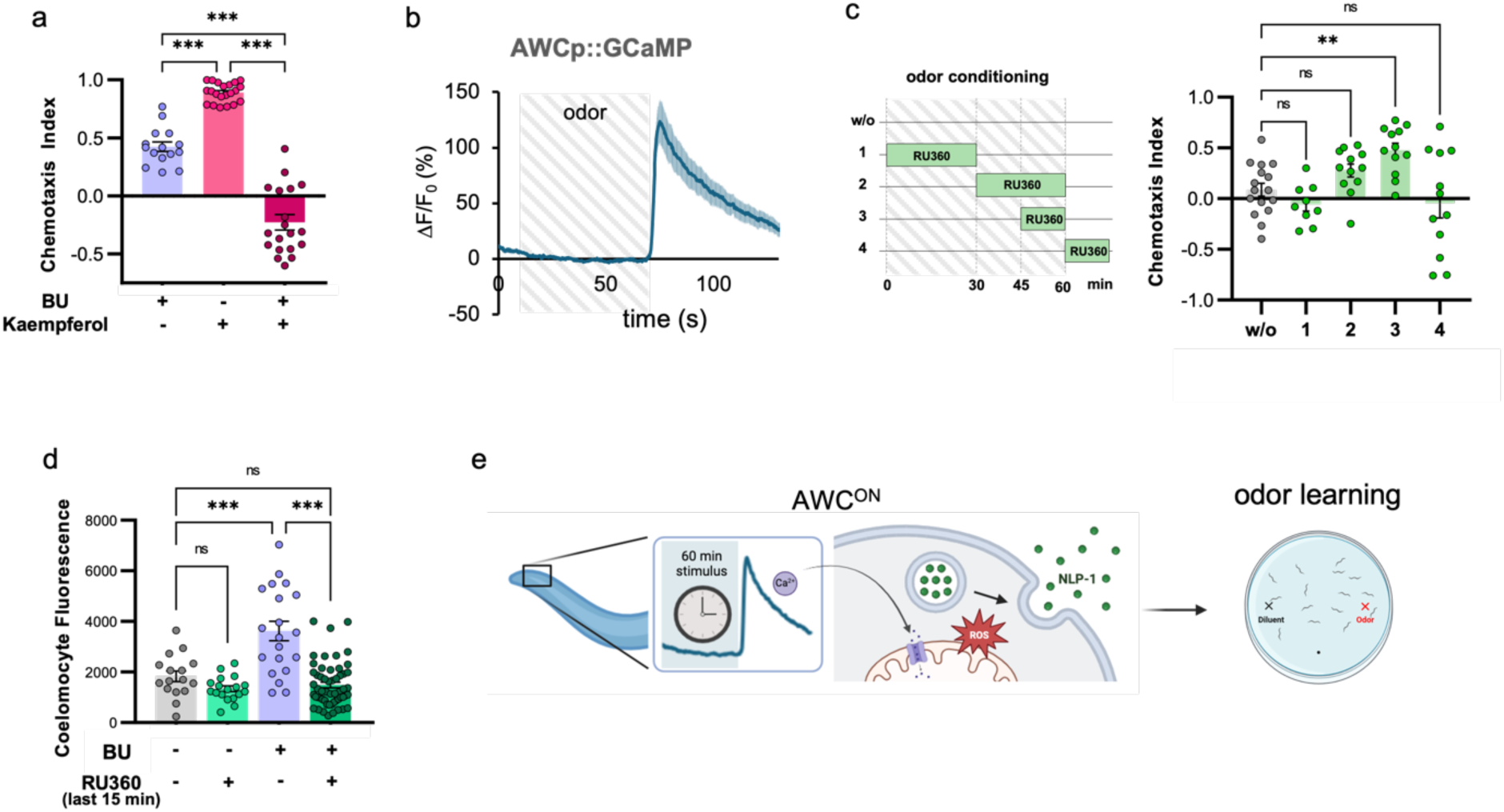
Coincident mtROS production and odor exposure is sufficient for odor learning. (A) Kaempferol treatment during 10 min odor conditioning results in strong learning. (B) Calcium response to odor addition and removal in the AWC neuron (n=12). (C) Effect of RU360 treatment at different time points during the 60 min butanone conditioning on odor learning. (D) NLP-1 secretion in worms treated with RU360 during the last 15 min of 60 min butanone conditioning. (E) Model of MCU activation in response to prolonged odor stimulus, which results in NLP-1 secretion and odor learning. P values were determined using one-way ANOVA with Dunnett’s multiple comparisons test. Asterisks indicate p value (ns, not significant; **p<0.01; ***p<0.001).

### MCU-1 function is required at the end of conditioning

Finally, we asked at what point during odor conditioning MCU is activated. This was because NLP-1 release is observed only after 60 min odor stimulus but not after 30 min (Fig 3D). How does the neuron respond differently to 30-min versus 60-min conditioning? Is the neuropeptide secreted in small quantities throughout the duration of conditioning, accumulating to a critical level that leads to learning, or is it released all at one time only after 60-min odor stimulus? One thing to consider is that the AWC sensory neuron is an “odor-OFF” neuron – it is inhibited by odor stimulus, and activated when the odor is removed(Chalasani *et al*, 2007) (Figure 5B). This implies that whether an odor is given 30 min or 60 min, there is only one calcium transient occurring at the end when the odor is finally removed. Since synaptic activity is usually associated with an active neuron, we hypothesized that MCU activation, and thus NLP-1 secretion, also occurs at the end of the stimulus.

To test this, we used RU360 to block MCU at specific time points during odor conditioning and tested their behavior (Fig 5C). Adding and removing MCU inhibitor RU360 in the first or last half of pre-exposure showed that blocking MCU during the second half of odor conditioning tended to suppress aversive learning (Fig 5C). Furthermore, blocking MCU function during the last 15 min was sufficient to prevent odor learning. This suggested that MCU function was required at the end of odor exposure, which coincides with the timing of the calcium transient that appears upon odor removal. When RU360 was added during the washing step immediately after odor exposure, it had no effect on odor learning (Fig 5C). This was not surprising, as there is likely a significant time delay for the inhibitor to reach the neuron.

Finally, we tested whether the adding RU360 at the end of odor conditioning prevented NLP-1::Venus secretion. Consistent with their chemotaxis behavior, worms that were supplemented with RU360 during the last 15 min of odor conditioning failed to show NLP-1::Venus accumulation in the coelomocytes (Fig 5D). These results strongly suggest that MCU-mediated calcium influx, as well as NLP-1 secretion, takes place at the end of 60 min conditioning, likely at the time of AWC calcium transient upon odor removal (Fig 5E). Since calcium transient occurs whether after 30 min or 60 min odor stimulus, this hints at a mechanism to selectively activate MCU in response to prolonged odor conditioning.

## DISCUSSION

Mitochondrial dysfunction is known to accompany many psychiatric, neurological, and neurodegenerative disorders. However, mechanistic studies on how the dysfunctions contribute to these disorders are lacking. Our findings identify a specific aspect of mitochondria function – control of calcium influx and mtROS production – that contributes to normal brain function and behavior.

In this study, we showed that MCU-1, and thus mitochondrial calcium, contributes to behavioral plasticity in *C. elegans* by regulating neuropeptide release at the sensory neuron. We found that *mcu-*1 mutants are defective for 60 min aversive odor learning, and this was due to lack of NLP-1 secretion that normally occur after 60 min odor conditioning. Odor learning and NLP-1 secretion required MCU activity and mtROS production in the sensory neuron at the time of conditioning. Correspondingly, ectopic MCU activation or mtROS during odor exposure accelerated learning. Furthermore, we showed evidence that MCU activation and NLP-1 secretion occur at the same time as the AWC calcium transient, which hints at a mechanism to selectively activate MCU in response to prolonged odor conditioning. The study shows that mitochondria in neurons play a direct role in determining the composition of synaptic signals during neuronal activity – whether to release a specific neuropeptide in addition to the classical neurotransmitter. The release of this neuropeptide alters the animal’s behavior to display the appropriate level of response to the experienced odor.

A previous study in Drosophila also reported odor memory defects in MCU mutants (Drago & Davis, 2016). However, in flies, MCU was required for the proper development of the mushroom body neurons, and thus silencing MCU after pupation had no effect on memory. This demonstrates the many roles mitochondria and mitochondrial calcium play in different systems and organisms, and highlight the benefit of studying gene function in multiple model organisms.

### MCU responds to conditioning duration

Since its molecular identity was discovered, studies have identified various regulatory mechanisms of MCU function. In addition to the pore-forming subunit MCU-1, the MCU complex includes auxiliary subunits MICU and EMRE that regulate channel activity. MICU senses calcium through its EF-hand domain, causing MCU to open in response to cytoplasmic calcium increase(Vais *et al*, 2020). Consistently, simultaneous imaging of cytoplasmic and mitochondrial calcium in neurons shows mitochondrial calcium influx occurring concurrently with cytoplasmic calcium increase(Katsenelson *et al*, 2022). In addition to MICU and EMRE, additional factors were found to regulate MCU activity in mice studies, such as CAMKII, MCUR1, or mitochondrially localized IGFR, suggesting that MCU responds to multiple cues in addition to cytoplasmic calcium concentration(Katsenelson *et al*., 2022; Lin *et al*, 2019; Vais *et al*, 2015). This is consistent with the emerging view that MCU-mediated calcium influx is selective, occurring in response to more significant changes in cellular activity. Multiple studies show that mitochondrial calcium influx is usually coupled to either stronger electrical stimulation of the cell or localized cytoplasmic calcium increase(Alvarez-Illera *et al*, 2020; Csordas *et al*, 2013; Doser *et al*, 2023; Groten & MacVicar, 2022; Katsenelson *et al*., 2022; Lin *et al*., 2019; Stoler *et al*, 2022).

Our results indicate that MCU activity in the AWC neuron represents a threshold for the difference between 30 min and 60 min aversive odor learning. This suggests that MCU, and hence mitochondrial calcium entry, may act as a duration sensor for sensory conditioning (Fig 6F). How may this occur? Contrary to most other sensory stimuli, extended odor stimulation in the AWC is characterized by prolonged inactivity rather than activity. However, a careful characterization of AWC calcium response has previously shown that progressively longer odor exposure time from 10 s, 1 min, to 3 min, resulted in a progressively faster and higher calcium spike at the synaptic terminal upon odor removal(Ventimiglia & Bargmann, 2017). This shows that cytoplasmic calcium can potentially reflect the duration of odor stimulation. Whether this also applies to 30 min and 60 min odor exposure remains to be seen, but it is one possible way that different durations of odor stimulation could be signaled in the cell.

In *C. elegans* neurons, it was previously shown that mitochondrial H_2_O_2_ increases the release of the neuropeptide FLP-1 from the AIY neuron, inducing the oxidative stress response in peripheral tissues(Jia & Sieburth, 2021). The study showed that different bacterial diets, thus internal metabolic cues, can cause mitochondrial calcium influx, as well as mtROS and H_2_O_2_ production, from the AIY axonal mitochondria. Our study adds to the evidence that this mechanism of neuropeptide secretion may be a general process that occurs in other neurons in *C. elegans*, and in response to a wider variety of cues beyond metabolic, such as odor stimuli. We note, however, that internal metabolic cues may still be involved in this case, as aversive odor conditioning requires food deprivation. Past studies show that insulin-like signaling mediates the response to starvation and that this is required for aversive odor learning (Cho *et al*, 2016; Lin *et al*, 2010). If so, this suggests the possibility of AWC mitochondria acting as an intersectional hub to integrate sensory information and internal metabolic states.

### mtROS and neuropeptide release

Mitochondrial calcium influx is a prerequisite for hormone secretion in endocrine cells, most notably insulin from pancreatic β cells(Quan *et al*, 2015). In β cells, glucose uptake and subsequent metabolic processes trigger mitochondrial calcium influx, as part of a process known as metabolism-secretion coupling(Maechler *et al*, 1997; Wiederkehr *et al*, 2011). Mitochondrial calcium influx produces “coupling factors” that together with other signals contribute to insulin release, among which include mtROS and H_2_O_2_(Leloup *et al*, 2009; Pi *et al*, 2007). Similar to our study, treatment with antioxidants prevented insulin release, showing that hormone secretion and neuropeptide secretion share similar mechanisms.

Neurons often express several neuropeptides but whether they are co-released or controlled individually is not well understood. Single-cell sequencing technology is beginning to reveal the prevalence of the existence of multiple neuropeptides per single neurons(Hanchate *et al*, 2020; Smith *et al*, 2019). This raises the question of what kinds of mechanisms exist that would allow for separate regulation of neuropeptide release from the same neuron(Klose *et al*, 2021). Our study suggests one possible mechanism of differential release. We show that the release of another highly expressed neuropeptide, NLP-3, does not respond to MCU activation. Similar selectivity in neuropeptide release was also observed in the AIY interneuron of *C. elegans*, where mtROS caused the release of the FMRFamide-like neuropeptide FLP-1 but not FLP-18(Jia & Sieburth, 2021). What determines whether a neuropeptide is controlled by the mitochondria will be a topic with important implications.

### Neuronal mitochondria and behavior

Mitochondria carry out a range of functions, from energy production, calcium buffering, lipid biosynthesis, ROS signaling, to signaling cell death. In cells with extreme structural polarity such as neurons, these range of functions often serve many compartment-specific roles, adding to the complexity of their function. Understanding the varied roles of mitochondria in neurons will help illuminate its contribution to various psychiatric, neurological, and neurodegenerative diseases. One study that directly linked mitochondrial function to behavior showed that reduced function and altered structure of mitochondria in the nucleus accumbens (NAc) result in reduced social dominance in mice (Hollis *et al*, 2015). Conversely, infusion of NAD^+^ precursor to improve mitochondrial respiration abolished the social disadvantage(Hollis *et al*., 2015).

Moreover, administering the anxiolytic diazepam also reverted the social disadvantage, by boosting NAc mitochondrial function through the release of dopamine into the NAc (van der Kooij *et al*, 2018). Finally, this effect could be blocked by blocking complex I function in the NAc. With the advent of improved tools to monitor brain activity and mitochondrial function, similar mechanistic studies exploring the causative links between mitochondria and behavior are expected.

## MATERIALS AND METHODS

### Worm cultivation

C. elegans were maintained at 20°C on plates containing nematode growth media (NGM) with OP50 bacteria. Unless otherwise specified, day 1 adults were used for experiments. All strains used in this study are listed in Table S1.

### Construction of plasmids and transgenic strains

Whole worm RNA was isolated using Direct-zol RNA Microprep kit (Zymo Research). cDNA was prepared using SuperScript IV First Strand Synthesis System (Invitrogen). *mcu-1* sequence was amplified from worm cDNA using PrimeSTAR MAX DNA Polymerase (Takara) and cloned into pCR-Blunt II-TOPO vector (Invitrogen). This was used to amplify *mcu-1* with the appropriate restriction enzyme sites required for the different plasmids. roGFP sequence was amplified from genomic DNA of KWN119 strain (gift from Nehrke lab) which expresses mito-roGFP under the *myo-3* muscle-specific promoter. Primers that bind to the 3’ end of the *myo-3* promoter and reverse primer binding to the *unc-54* 3’UTR sequence were used to amplify the whole fragment, then existing enzyme restriction enzyme sites were used to cut the insert, which was ligated into the AWC-specific *ceh-36p* plasmid (gift from Juang lab). MiniSOG sequence was amplified from genomic DNA of CZ20310 strain (CGC), which expresses HIS-72 tagged miniSOG in the germline. The first 228 nt sequence of *tomm-20,* which includes 165 nt coding sequence and the first intron, was amplified from worm genomic DNA. Venus sequence was cloned from a worm plasmid containing the sequence (gift from Sieburth lab). *nlp-1*, and *nlp-3* sequence was amplified from worm cDNA. All primers used in this study are listed in Table S2.

To generate transgenic worm strains, plasmids were injected at 20 ng/μl, along with 40 ng/μl *unc-122p::gfp* or *mCherry* co-injection marker plasmids. pUC19 plasmid was used to bring the total DNA concentration of the injection mix to 100 ng/μl.

### Chemotaxis assay

Assay was performed as described previously(Colbert & Bargmann, 1995). For assay plates, 10 ml of agar media (1.6% Difco Granulated Agar (Beckton Dickinson, Franklin Lakes, NJ, USA), 1 mM CaCl_2_, 1 mM MgSO_4_, 5 mM KPO_4_) was poured into 10 cm petri dish and were left out to dry for at least 1 day. Well-fed, day 1 adult worms were used for all assays. Odors were diluted as follows unless otherwise stated: 1 μl Benzaldehyde (Sigma) in 200 μl ethanol, 1 μl 2-Butanone (Sigma) in 1 ml ethanol, 1 μl Diacetyl (Waco) in 5 ml, and 1 μl 2,3-Pentanedione (Alfa aesar) in 10 ml ethanol. For extrachromosomal transgenic strains, we scored only the transgenic strains that expressed the fluorescent co-injection marker.

### Aversive associative learning assay

Day 1 adults were collected from plates and washed three times with S-basal buffer. For conditioning, worms were placed in 1 ml of S-basal buffer with or without each odorant with the following concentrations: Benzaldehyde 1:10,000, 2-Butanone 1:10,000, Diacetyl 1:5,000, 2,3-Pentanedione 1:100,000. The microcentrifuge tubes were rotated on a rotator for conditioning, then washed twice with s-basal buffer and once with 3DW before conducting the chemotaxis test. For heat-shock-inducible *mcu-1* mutant worms, synchronized worms at the L1 stage were transferred to seeded NGM plates to grow until ready for heat shock. At each developmental stage, plates of worms were sealed with parafilm and immersed into a 32°C water bath for 1 hr. Plates were moved back to 20°C until day 1 stage when the aversive learning assay was conducted. Worms heat-shocked at the adult stage were placed at 20°C for 2 hr until used for the assay. For mito-miniSOG strains, worms were grown in plates covered with aluminum foil to prevent excess light exposure during development. We found that for learning and NLP-1 secretion, these strains were sensitive to ambient light and this was sufficient to induce accelerated odor learning and NLP-1::Venus secretion (below). Thus, odor conditioning and subsequent chemotaxis assays were conducted either on the bench (light+) or in a dark room to avoid ROS generation by ambient light (light-).

### Pharmacological treatment

All pharmacological substances used in this study, except for juglone and H_2_O_2_, necessitated permeabilizing the cuticle. This was accomplished by growing worms in *dpy-8* RNAi bacteria at the last molting stage, from L4 to adult stage. *dpy-8* RNAi bacteria clone was obtained from the Ahringer RNAi library(Kamath *et al*, 2003) (Source BioScience). RNAi plates were prepared as previously described(Ahringer). Permeabilization of cuticle did not affect their behavior (data not shown). Following are the stock solutions used: 6.358 mM Ruthenium Red in water (Millipore), 0.5 mg/mL RU360 (Millipore) in deoxygenated water, 50 mM of Kaempferol (Santa Cruz Biotechnology) in EtOH, 50 mM of Juglone (Sigma-Aldrich) in DMSO, 100 mM of N-Acetyl-L-Cysteine (BioBasic) in water, 18.938 mM of mitoTEMPO (Santa Cruz Biotechnology) in DMSO, and 30% H_2_O_2_ (Sigma-Aldrich). For behavior assay, about 1000 adults worms were transferred into a 1.5 mL Eppendorf tube with S-basal, and washed three times. Pharmacological treatments were added to washed worms at final concentration of 100 μM for Ruthenium Red, 0.5 μM for RU360, 100 μM for Kaempferol, 300 μM for Juglone, 0.5 mM for N-Acetyl-L-Cysteine, 10 μM for mitoTEMPO, and 5 mM H_2_O_2_.

### Microscopy and analysis

All images were obtained using Zeiss LSM800 confocal microscope with Zeiss C-Apochromat 40x/1.2 W Korr objective and ZEN 2.6 software. For coelomocyte imaging, 30–40 L4 stage animals were collected from NGM plates, washed three times, and exposed to either butanone (1:10,000 in S-basal) for 1 hr or to each pharmacological treatment for 10 min in S-basal. After treatment, worms were briefly washed, then paralyzed using sodium azide and mounted on 2% agarose pads. All coelomocytes along the length of the worm was imaged. Maximum intensity projection was obtained using image stacks and used for analysis. Fluorescence was quantified by manually delineating each coelomocyte using the polygon selection tool and measuring the integrated density in ImageJ.

For the mito-miniSOG strain, we found that even ambient light was enough to significantly increase NLP-1::Venus in the coelomocytes. Worms were grown in plates wrapped in foil to prevent excess light exposure, but light exposure was inevitable during microscopic analyses. We also confirmed that, although ambient light produces ROS, only strong blue light (30 min) was able to induce AWC ablation (Figure S2).

To image mito-roGFP and calculate reduction/oxidation ratios, worms were imaged at two different wavelengths for excitation, 488/405 for the reduced/oxidized (R/O) ratio(Vevea *et al*, 2013). Maximum intensity projection was obtained for each image. After setting the ROI for each mitochondrion in image J, the fluorescence intensity of the reduced mitochondria was measured, and the same ROI is applied to the oxidized mitochondria to obtain the ratio of 2 stacks after subtracting the background value from the two values. To observe the range of roGFP redox ratios, 5 mM H_2_O_2_ for 10 min was used.

### Calcium Imaging and analysis

Worms were immobilized in a poly(dimethylsioloxane) (PDMS) microfluidic chamber described previously(Chronis *et al*, 2007). Using the microfluidic chamber design generously provided by the Bargmann lab, the SU-8 micro mold and PDMS chip were constructed by MicroFit (Hanam, Republic of Korea). Day 1 adult worms were transferred to unseeded plates to clean bacteria off the body, then transported to the chamber in S-basal buffer via polyethylene tubing (Intramedic). Benzaldehyde at 1:100,000 dilution was used for imaging. GCaMP fluorescence was visualized at 40x magnification using IX-73 inverted fluorescent microscope (Olympus) attached to Iris 9 sCMOS camera (Photometrics) and a pE-800 LED illuminator (CoolLED). Fluorescence was recorded using MetaFluor 3.1 at approximately 5 frames/s. Resulting image sequence was processed using the Fiji software (NIH). ROI was aligned using the ‘align slices in stack’ function of the Template Matching plugin, then pixel intensity of AWC cell body was measured.

### Statical analysis

All data are expressed as means ± standard error of the mean (SEM). Statistical significance was determined by student’s t test or one-way ANOVA. Asterisks indicate: *p<0.05, **p<0.01, ***p<0.001, and ****p<0.0001. All experiments were performed in at least 3 independent trials on different dates.

## ACKNOWLEDGEMENTS

We thank the Caenorhabditis Genetics Center (CGC) (P40 OD010440) and the National BioResource Project (NBRP) for providing several strains used in this study. Venus plasmid was generously shared by Derek Sieburth. Worm strain expressing myo-3p::mito-roGFP (KWN119) used to clone mito-roGFP was shared by the Nehrke lab. Worm expression plasmids containing the *ceh-36* promoter and *hsp-16.2p* promoter were gifted by Bi-Tzen Juang. Kyuhyung Kim and Wonchan Choi provided valuable assistance with setting up calcium imaging. Layout for the microfluidic chip design used for calcium imaging was provided by the Bargmann lab. We thank Jin Lee, Kyung Won Kim and members of the Kim lab, as well as members of OMRC for helpful discussions and comments on the manuscript. This research was sponsored by the National Research Foundation of Korea 2017R1A5A2015369, RS-2024-00409403 (K.-S.P.), 2019R1C1C1008708 (K.-h.Y.), and the Ministry of Science and ICT, Bio and Medical Technology Development Program 2020M3A9D8039920 (K.-S.P).

## Author contributions

Conceptualization, H.K.L., K.S.P., and K.-h.Y.; Investigation, H.K.L., D.K.J and K.-h.Y.; Writing – Original Draft, H.K.L. and K.-h.Y.; Writing – Review & Editing, H.K.L., K.S.P., and K.-h.Y.; Funding Acquisition, K.S.P. and K.-h.Y.; Supervision, K.S.P. and K.-h.Y.

---

**Figure S1.**
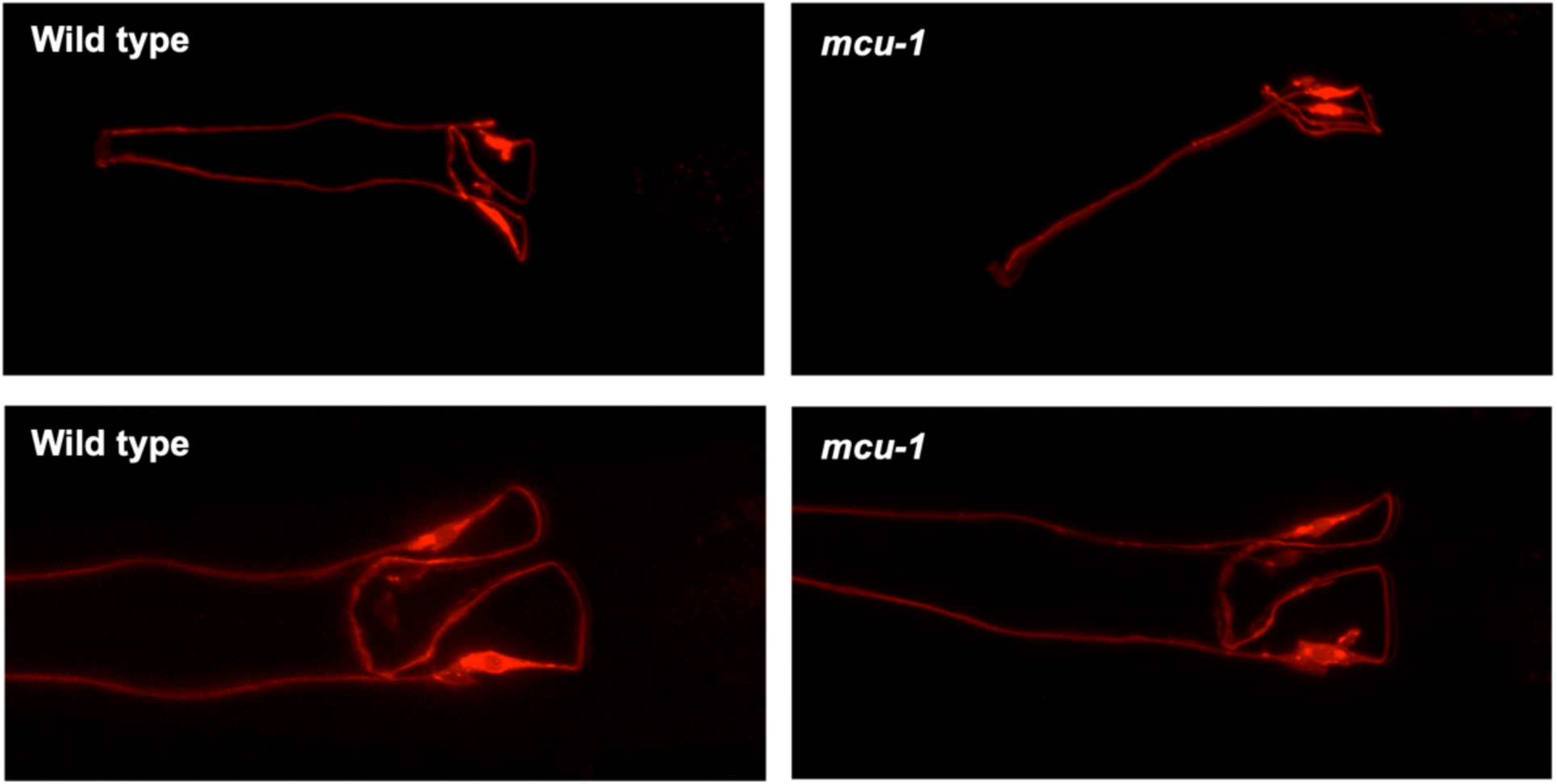
AWC morphology in wild type and mcu-1 mutant worms.

**Figure S2.**
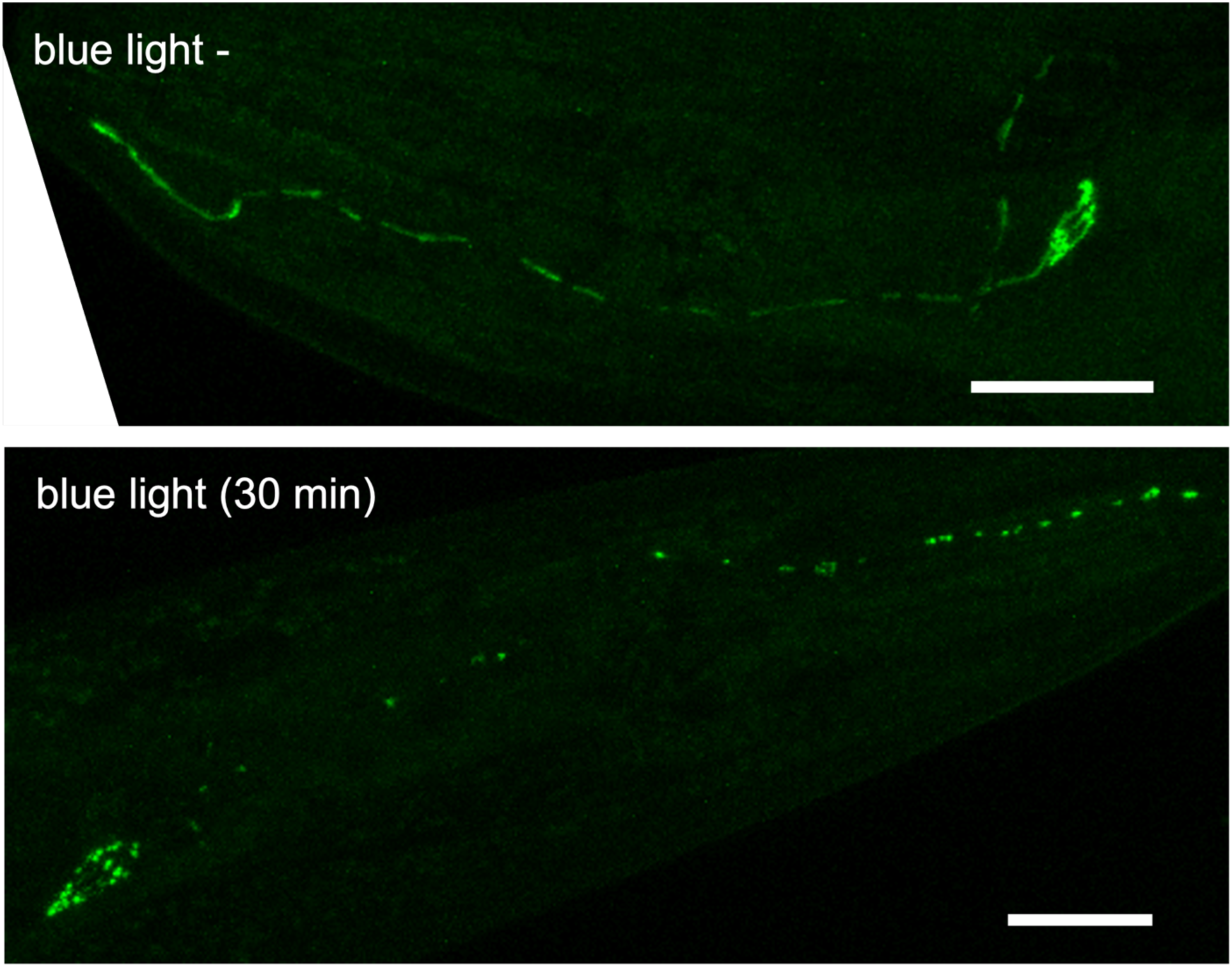
Strong blue light is required for mito-miniSOG to cause mitochondrial loss and fragmentation. GFP labeling of mitochondria in the AWC neuron show that exposure to ambient light is not sufficient to cause cell ablation (blue light -). Irradiation with intense blue light for 30 min resulted in severe loss and fragmentation of mitochondria when observed immediately after the treatment (blue light +). Scale bars indicate 20 μm.

## Notes

### Competing Interest Statement

The authors have declared no competing interest.

### Summary of Updates

This updated version of the manuscript includes additional data to better suport our claims that mitochondrial calcium influx occur selectively to longer sensory input in the AWC neuron. In addition, the overall focus of the manuscript has been modified to emphasize the significance of MCU's role in natural behavior and learning.

