## Supplementary material for "Mitochondrial calcium modulates odor-mediated behavioral plasticity in *C. elegans*": Table S1, S2

**Table S1.** All strains used for this study

| Experimental Models: Organisms/strains | Source | Identifier |
|---|---|---|
| N2 | CGC | N2 |
| *mcu-1 (tm5026)* | NRBP | |
| *mcu-1 (tm5026); okyEx101[rab-3p::mcu-1 + unc-122p::gfp]* | This study | KHY010 |
| *mcu-1 (tm5026); okyEx103[ceh-36p::mcu-1 + unc-122p:mCherry]* | This study | KHY129 |
| *mcu-1 (tm5026); okyIs100[hsp-16.2p::mcu-1 + unc-122p::mCherry]* | This study | KHY231 |
| *okyEx131[ceh-36p::mito-roGFP + unc-122p::mCherry]* | This study | KHY170 |
| *mcu-1 (tm5026); okyEx131[ceh-36p::mito-roGFP + unc-122p::mCherry]* | This study | KHY266 |
| *okyEx132[ceh-36p::nlp-1::Venus + unc-122p::mCherry]* | This study | KHY220 |
| *mcu-1 (tm5026); okyEx132[ceh-36p::nlp-1::Venus + unc-122p::mCherry]* | This study | KHY221 |
| *okyEx147[ceh-36p::tomm-20::miniSOG::SL2::mCherry]* | This study | KHY263 |
| *okyEx132[ceh-36p::nlp-1::Venus + unc-122p::mCherry], okyEx147[ceh-36p::tomm-20::miniSOG::SL2::mCherry]* | This study | KHY264 |
| *okyEx134[ceh-36p::nlp-3::Venus]* | This study | KHY265 |
| *nlp-1(ok1470)* | This study | RB1341 |
| *IskEx1553[ceh-36-Δ1p::GCaMP3 + unc-122p::dsRed]* | Gift from Dr. Kyuhyung Kim | KHK21690 |

**Table S2.** All primers used for this study.

| **Cloning Primers** | **Forward 5'-3'** | **Reverse 5'-3'** |
|---|---|---|
| Prab-3 | tcgtcgCTGCAGATCTTCAGATGGGAGCAGTGGAC | tgatgaGGATCCTGCTTTTTTGTACAAACTTGTCATC |
| mcu-1 | ATGAGGAATGGCCGATGCT | TTACTTTTCAGCTTCCAAATTGGAT<br>attattGCTAGCTTACTTTTCAGCTTCCAAATTGGAT |
| Venus | aataGGTACCAGTAAAGGAGAAGAACTTTTCACTGG | tattGAATTCTTATTTGTATAGTTCATCCATGCCATGT |
| nlp-1 | aataGGATCCACATCAACTTGAGGCAACGATGA | aataGGTACCACGACGTCCCAATCCGACAAAG |
| nlp-3 | aataGGATCCATGAGCAAAATCGTCGCTTGC | aataGGTACCATAGTAATTTTCCAACATTTCATATCGATTG |
| miniSOG | aaatACCGGTGAGAAAAGTTTCGTGATAACTGATCCA | aaatGAATTCAAATGGTACCTTATCCATCCAGCTGCACTCC |
| tomm-20 mitochondria targeting sequence | attattGGATCCATGTCGGACACAATTCTTGGTTTC | aaatACCGGTTGCTCCAGCCTGGGCACGTC |

| | | |
|---|---|---|
| SL2 | aatGGTACCCGCTGTCTCATCCTACTTTCACC | ataatCTCGAGAGCAGTTTCCCTGAATTAAAATTAGAAG |
